# *In-Vitro* Efficacy of Targeted FODMAP Enzymatic Digestion (FODZYME®) in a High-Fidelity Simulated Gastrointestinal Environment

**DOI:** 10.1101/2022.05.16.492152

**Authors:** Kenny Castro Ochoa, Shalaka Samant, Anjie Liu, Cindy Duysburgh, Massimo Marzorati, Prashant Singh, David Hachuel, William Chey, Thomas Wallach

**Affiliations:** SUNY Downstate Health Sciences University, Division of Pediatric Gastroenterology. Brooklyn, NY; Kiwi Biosciences. Cambridge, MA; Prodigest. Ghent, Belgium; University of Michigan, Division of Gastroenterology. Ann Arbor, Michigan

**Keywords:** IBS, FODMAP, Fructan, Generally-Regarded-as-Safe

## Abstract

**Introduction:** Irritable bowel syndrome (IBS) is characterized by abdominal pain and changes in bowel habits. FODMAPs are poorly absorbed short-chain carbohydrates that may drive commensal microbial gas production, promoting abdominal pain in IBS. Low-FODMAP diet can result in symptomatic improvement in 50-80% of IBS patients. However, this diet is not meant to be sustained long term, with concern for downstream nutrition and microbial issues. In this study, we evaluate the function of a targeted FODMAP enzymatic digestion food supplement FODZYME® containing an fructan hydrolase enzyme in a simulated gastrointestinal environment.

**Methods:** Using SHIME®, a multi-compartment simulator of the human gut, FODZYME® dose finding assay in modeled gastrointestinal conditions assessed enzymatic ability to hydrolyze 3 g of inulin. Full intestinal modeling assessing digestion of inulin, absorption of fructose, gas production and other measures of commensal microbial behavior was completed using 1.125 g of FODZYME®.

**Results:** After 30 minutes, 90% of the inulin was converted to fructose by 1.125 g of FODZYME®.Doubling dosage showed no significant improvement in conversion, whereas a half dose decreased performance to 77.2%. 70% of released fructose was absorbed during simulated small intestinal transit, with a corresponding decrease in microbial gas production, and a small decrease in butyrate and short chain fatty acid (SCFA) production.

**Discussion:** FODZYME® specifically breaks down inulin in representative gastrointestinal conditions, resulting in decreased gas production while substantially preserving SCFA and butyrate production in the model colon. Our results suggest dietary supplementation with FODZYME® would decrease intestinal FODMAP burden and gas production.

## Introduction

Irritable bowel syndrome (IBS), a disease of gut-brain interaction (DGBI) characterized by abdominal pain, bloating, and changes in bowel habits (constipation, diarrhea, or both) not explained by another medical etiology, impacts up to 15% of the population of the United States^1,2^. Up to 84% of patients with IBS report food as a key trigger for their gastrointestinal symptoms^3^. In particular, fermentable oligosaccharides, disaccharides, monosaccharides, and polyols (FODMAPs) have been shown to induce gastrointestinal symptoms in IBS patients in a dose-dependent manner^4-6^. FODMAPs are non-absorbable, osmotic carbohydrates that undergo bacterial fermentation in the colon. The effect of FODMAPs on IBS symptoms appears to be in part driven by the luminal distention caused by fermentation products such as hydrogen and methane^7^. However, change in the intestinal microenvironment due to microbial metabolites or acid production also likely plays a role in FODMAP-mediated IBS pathophysiology^8^.

Given the contribution of FODMAPs towards IBS symptoms, several clinical trials have evaluated the role of a low FODMAP diet in the management of IBS and consistently shown symptom improvement in 50-80% of the IBS patients ^5, 9, 10^. Clinical implementation of the low FODMAP diet comprises three phases to be followed by IBS patients: i. ‘restriction’ involves dietary exclusion of FODMAPs for 4–6 weeks to test for response to the diet ii. ‘reintroduction’, which helps to identify the impact of specific FODMAPs and their doses on symptoms, iii. ‘personalization’, involves development of a long-term plan that can achieve dietary variety coupled with symptom control^11^. A recent network meta-analysis pooled data from 13 randomized controlled trials evaluating the efficacy of a low FODMAP diet in IBS and found that the low FODMAP diet was superior to other dietary interventions in achieving improvement in global IBS symptoms, abdominal pain, and bloating^5^. Despite the clinical efficacy of a low FODMAP diet in IBS, some potentially unfavorable consequences have been reported^12^. The low FODMAP diet is a complex intervention and requires specialist and formal dietetic input prior to implementation which can impose substantial pressures on publicly-funded healthcare services^13^. Some other reported drawbacks of the low FODMAP diet are social isolation, difficulties with travel and eating out, and the additional cost of specialty food items to comply with the dietary requirements without introducing high FODMAP foods^14^. Thus, implementing a low FODMAP diet is cumbersome, restrictive, time-consuming, and costly. Furthermore, a few studies suggest that the restriction phase of a low FODMAP diet may be associated with reduced dietary intake of some micronutrients (e.g., iron and thiamine) and may lead to a reduced fecal abundance of short-chain fatty acids (derived from bacterial fermentation of FODMAPs) and putatively beneficial bacteria such as Bifidobacteria^8, 9, 15, 16^. Moreover, a long term study of IBS patients following the low FODMAP diet has also reported reduced intake of certain nutrients, and reduced abundance of short-chain fatty acids and beneficial bacteria, all of which can affect colonic health^17^.

Given the challenges of the low-FODMAP diet, alternative approaches that may decrease gas production secondary to bacterial fermentation of FODMAPs without fully removing fermentable carbohydrates may prove more feasible for patient use, as well as more conducive to long-term implementation. To achieve this goal one possible avenue of intervention could include the use of enzymatic digestion of common FODMAPs, decreasing their availability to the intestinal flora without fully removing the benefits of fermentable fiber for colonic health. This approach would also facilitate the continued intake of other beneficial nutrients from FODMAP-rich foods. Enzymes have previously been successfully applied to breaking down specific non-absorbable oligosaccharides before they are metabolized by colonic bacteria leading to reduced abdominal pain, bloating, flatulence and diarrhea^18^. Some nonprescription dietary supplements with specific FODMAP hydrolyzing enzymes such as lactase (*e*.*g*. Lactaid® or Lactrase®) and α-galactosidase (*e*.*g*. Beano® or Bean Relief™) are commercially available for managing FODMAP related symptoms. Application of FODMAP degrading enzymes to food raw materials and during food processing is also being explored^19^.

Emerging data suggests that fructans are the most consistent FODMAP subgroup to trigger symptoms in IBS patients sensitive to FODMAPs^20, 21^. Fructans are very commonly ingested FODMAP carbohydrates found in several vegetables such as onion, garlic, and leak, some fruits such as bananas, and grains such as wheat and rye that are consumed in large amounts^7, 19^. They are linear or branched, oligomers or polymers consisting primarily of fructosyl-fructose units with or without one glucosyl unit^22^. The most common structural form of fructans present in everyday foods are inulin-type fructans^23, 24^. Since humans lack the enzymes hydrolyzing fructans to fructose, these polymers cannot be absorbed in the intestine. The malabsorbed fructans are delivered to the colon, where they are fermented by the resident bacteria leading to gas production, which accompanied by the fructan molecules drawing water into the intestine causes luminal distention which can exacerbate the symptoms of IBS^25^. Moreover, a double-blind, placebo-controlled RCT also found fructans as the likely culprit for gastrointestinal symptoms in patients with non-celiac wheat sensitivity^20^. FODZYME®, is a proprietary combination of generally regarded as safe (GRAS) enzymes designed for application to meals, just before ingestion, with the goal of pre-digesting fructans before they reach the colon. In this study, we evaluate the function of FODZYME®, in a simulated gastrointestinal environment to determine its efficacy in common gastric conditions such as IBS.

## Materials and Methods

### SHIME® model gastrointestinal tract

The reactor setup was adapted from the Simulator of the Human Intestinal Microbial Ecosystem (SHIME®), representing the gastrointestinal tract (GIT) of the adult human, as described by de Wiele *et al*^26^. The SHIME® consists of a succession of five reactors simulating the different parts of the human gastrointestinal tract. The first two reactors are of the fill-and-draw principle to simulate different steps in food uptake and digestion, with peristaltic pumps adding a defined amount of SHIME® feed and pancreatic and bile liquid, respectively, to the stomach and small intestine compartment and emptying the respective reactors after specified intervals. The last three compartments – continuously stirred reactors with constant volume and pH control – simulate the ascending, transverse, and descending colon. Retention time and pH of the different vessels are chosen in order to resemble *in vivo* conditions in the different parts of the gastrointestinal tract. All vessels are kept anaerobic by flushing them with N2, and are continuously stirred and kept at 37°C.

### Dose-Finding Assay

This assay simulated gastric conditions in order to identify ideal dose- to-substrate ratios. Inulin, a linear polymer of β (2,1)-linked fructose residues, with terminal glucose residues found abundantly in wheat, onions, garlic and leeks was used as a representative substrate^27^ Inulin was obtained from Jarrow Formulas, USA. Typical SHIME® feed contains (in gm/liter) arabinogalactan (1.0), pectin (2.0), xylan (1.0), starch (4.0), glucose (0.4), yeast extract (3.0), peptone (3.0), mucin (1.0), and L-cysteine-HCl (0.5). A sugar-depleted gastric medium (i.e., without arabinogalactan, pectin, xylan, starch, and glucose) was applied in this study since inulin was added as a substrate at a concentration of 3000 mg/reactor. To closely mimic gastric conditions, phosphatidylcholine (0.17 mM) was added and salt levels were maintained at approximately 50 mM NaCl and approximately 7 mM KCl. Pepsin was excluded from the gastric medium in order to resemble a protease-free environment. Other aspects were completed as previously described^26^. Contents were incubated for 2h at 37°C while mixing via stirring. To simulate gastric pH shifts, a sigmoidal decrease in pH from 5.5-2.0 was induced. A blank control (without any enzyme), alternative carbohydrate-hydrolyzing enzyme control (‘dummy control’ - 0.225 g of alpha-galactosidase and 0.15 g of lactase), and three dosing levels (0.75, 1.125, and 1.875 g of FODZYME® enzymatic mixture, respectively) were compared. Fructose, the predominant breakdown product of inulin, was measured using high-performance anion-exchange chromatography with pulsed amperometric detection (HPAEC-PAD), at 30-, 60-, 90-, and 120-minute intervals from the point of enzyme addition. Degree of digestion (%) was determined for each of the test conditions at each time point by dividing the fructose content by the inulin concentration supplemented at the start of the incubation (i.e., 3000 mg/reactor). To determine the efficacy of FODYZME® in the presence of proteolytic activity encountered during gastric transit, this experimental set-up was repeated with the application of pepsin at 4000U/ml.

### Small Intestinal Modeling

To simulate a fed gastrointestinal state, contents derived from gastric digestion were moved through additional stages of the SHIME® apparatus. Here, while mixing via stirring, pH initially automatically increased from 2.0 to 5.5 within a period of 5 minutes, after which the pH of the medium increased from 5.5 to 6.5 during the first hour, 6.5 to 7.0 during the second hour, and finally remained constant at pH 7.0 during the third hour. The temperature was controlled at 37°C. A combination of a raw animal pancreatic extract (pancreatin) containing relevant enzymes in a specific ratio (5.6 TAME U/mL) as well as defined ratios of the different enzymes were used (i.e., 15.4 TAME U/mL for trypsin and 3.8 BTEE U/mL for chymotrypsin). Also, 10 mM bovine bile extract was supplemented to mimic bile salt action. In order to simulate the absorptive processes occurring in the small intestine, the duodenal phase (i.e., 30 min at pH 6.5) was followed by a dialysis approach (i.e., 3h at pH 7.0) using a cellulose membrane with a cut-off of 3.5 kDa. By introducing the small intestinal suspension within a dialysis membrane, molecules such as digested amino acids, sugars, micronutrients, and minerals were gradually removed from the upper gastrointestinal matrices. In order to reach an efficient absorption of small molecules, the dialysis solution (bicarbonate buffer) was refreshed every hour. HPAEC-PAD of luminal content at stomach end, duodenal end (30 minutes small intestinal exposure), and small intestinal end (end of exposure) were used to measure the fate of the fructose digestive product and determine fructose presentation to the simulated colon. The luminal fructose content represented bioaccessible fructose. To determine bioavailable fructose content, dialysate was sampled at 1, 2, and 3 hours after the start of dialysis and pooled for HPAEC-PAD. Degree of digestion (%) was determined for each of the test conditions at each time point by dividing the bioaccessible and bioavailable fructose content by the inulin concentration supplemented at the start of the upper GIT (i.e., 3000 mg/reactor).

### Colonic Modeling

Short-term single-stage colonic batch incubations were performed simulating the proximal colon environment. SHIME® colonic modeling uses a bacterial inoculum derived from a fresh fecal sample of a healthy adult male donor. At the start of the short-term colonic incubation, 20 ml of simulation medium from the upper GIT incubations was added to SHIME® nutritional medium containing basal nutrients that are present in the colon. As previously described, the carbohydrate-depleted colonic medium contained (in gm/liter) K_2_HPO_4_ (5.2), KH_2_PO_4_ (16.3), NaHCO_3_ (2.0), yeast extract (2.0), peptone (2.0), mucin (1.0), L-cysteine (0.5), and 2 ml /L Tween 80, pH 6.5^28^. Incubations were performed for 48 h, at 37°C under shaking (90 rpm) conditions. All experiments were performed in triplicate to account for biological variation. Several endpoints were measured. The degree of acidification during the experiment is a measure of the intensity of the bacterial metabolism (fermentation). The pH of the incubations was determined at 0h, 6h, 24h, and 48h after starting the incubation, thus giving an indication of the speed of fermentation. Gas production during the experiment is a measure of microbial activity and thus speed of fermentation of the potentially prebiotic substrates. Gas production was determined at 0h, 6h, 24h, and 48h after starting the incubation. Short-chain fatty acids (SCFAs) are an assessment of the microbial carbohydrate metabolism (acetate, propionate, and butyrate) or protein metabolism (branched SCFA) and can be compared to typical fermentation patterns for normal gastrointestinal microbiota. Samples for SCFA analysis were collected after 0h, 6h, 24h, and 48h of incubation. SCFA levels were measured using capillary gas chromatography coupled with flame ionization^29^.

### Statistical analysis

All experiments were performed in triplicate to account for variation. Prism GraphPad statistical software was used for all calculations. A confidence interval of 95% was applied (p<0.05), and significance calculations were performed using an unpaired t-test or Analysis of variance (ANOVA) as appropriate. If ANOVA was significant, posthoc analysis was performed for multiple comparisons using Tukey test.

### Ethics

Fecal samples were collected according to the ethical approval of the University Hospital, Ghent with reference number B670201836585.

## Results

### FODZYME® efficiently digests inulin in simulated gastric conditions

Low levels of fructose were measured in the blank control, indicating that low amounts of fructose were present in the inulin **[Figure 1A]**. Compared to blank control, the dummy control of alternative enzymes did not significantly alter the fructose content at any time point during the gastric incubation, suggesting no capacity of dummy enzymes to degrade inulin, indicating the need for specific enzymes to degrade the inulin substrate. On the contrary, FODZYME® was able to degrade the inulin substrate, as indicated by the detection of high fructose levels in FODZYME® replicated at all the measured time points **[Figure 1A]**. A significantly higher percentage of inulin digestion was noted at the end of gastric incubation (84.6-93.5%) for all three FODZYME® doses compared to blank or dummy control (P<0.0001 for each). Trials of three different dosages showed that inulin digestion percentage with standard dose of FODZYME® (1.125 g) was significantly higher than that seen with the FODZYME® dose of 0.75 g (P=0.0277), however, there was no significant difference between FODZYME® doses of 1.125 g and 1.875 g. Based on these findings, additional work went forward using the 1.125 g dosage.

**Figure 1:**
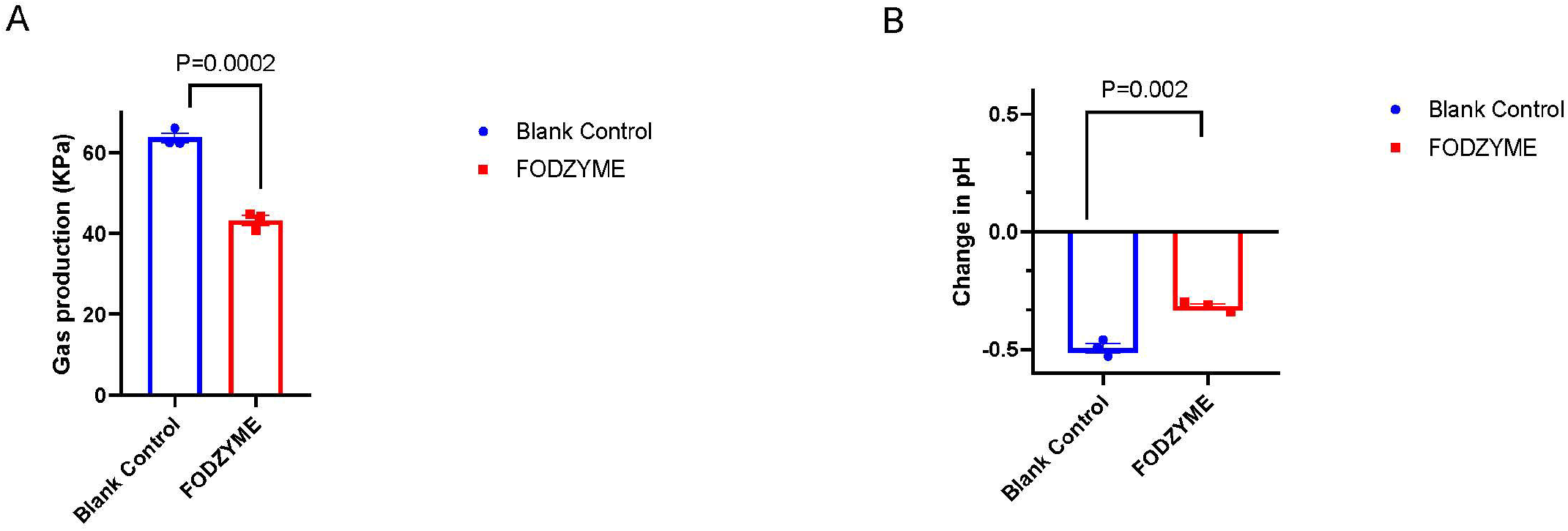
Simulated Gastric Survival of FODZYME®. A. Three dosing levels - 0.75 g (half dose), 1.125 g (standard dose), and 1.875 g (double dose) - of FODZYME® enzymatic blend were compared with blank control (no added enzyme) and dummy control (alternative carbohydrate - hydrolyzing enzyme), in the Simulator of the Human Intestinal Microbial Ecosystem (SHIME®), for their ability to degrade 3 g of inulin. Fructose, the predominant breakdown product of inulin, was measured using High-performance anion-exchange chromatography with pulsed amperometric detection (HPAEC-PAD), at 30-, 60-, 90-, and 120-minute intervals from the point of enzyme addition. Degree of digestion (fructose %) was determined for each of the test conditions at each time point by dividing the fructose content by the inulin concentration supplemented at the start of the incubation (i.e., 3000 mg/reactor). B. Same reaction and analysis conditions as A were repeated with the addition of pepsin (4000 U/ml). Data presented is for the 120-minute incubation interval. C. Following simulation of the absorptive processes that occur in the small intestine, HPAEC-PAD of luminal content at stomach end, duodenal end, and ileal end of the SHIME® were used to measure fructose amounts generated from inulin degradation following addition of FODZYME® vs a blank control (no added enzyme). Luminal fructose percentage was obtained by dividing the fructose content by the inulin concentration supplemented at the start of the incubation (i.e., 3000 mg/reactor). All experiments were performed in triplicate to account for variation. Error bars represent standard error.

### The enzymatic function of FODZYME® is only minimally affected by pepsin

A repeat study of FODZYME® performance in gastric conditions examined the impact of pepsin, a stomach protease that serves to digest proteins found in ingested food, on enzymatic function. In the presence of protease, we found a 21% decrease in the absolute digestive capacity of FODZYME® from 93.5% to 72.5% at the end of gastric incubation (p = 0.007) (**Figure 1B**). Overall, these experiments demonstrate the retention of significant enzymatic capacity, despite the presence of protease, in a high-fidelity gastric simulation. To mimic physiologic conditions, the rest of the experiments were performed in the presence of protease.

### Small intestinal modeling suggests fructose will be absorbed rather than presented to the Colon

As fructose itself is a FODMAP^22^, one concern in the use of a digestive enzyme is that it may not change the overall FODMAP burden in the intestine. SHIME® small intestinal modeling of control conditions^26^ vs. FODZYME® demonstrates substantial small intestinal absorption of digested fructose with only 16.2% of fructose remaining in luminal contents at the end of a small intestinal incubation of 3 hours **[Figure 1C]**. This result suggests that a significant portion of fructose generated from fructan degradation by FODZYME® can get absorbed into circulation, greatly reducing the FODMAP burden presented to the colon.

### Colonic modeling demonstrates a significant FODZYME® mediated reduction in bacterial gas, change in pH, with only moderate decreases in production of Butyrate and other short-chain fatty acids (SCFAs)

After small intestinal modeling, the contents were moved to a simulated colonic environment containing healthy donor fecal contents as per previously published protocol^26^. Overall, FODZYME® addition decreased gas production in the system by 16.71 kPa ((Control 63.67±2.1 kPa vs. FODZYME® 43.27 ± 2.2 kPa, P=0.0003) over 48 hours of colonic incubation **[Figure 2A]**. The bulk of gas production occurred over the first 6 hours, again with FODZYME® decreasing gas production from 49.70 ± 0.44 kPa in the control to 32.8 ± 3.4 kPa, for a 31.5% reduction in gas production (P=0.001). In our control sample, pH in the first 6 hours of exposure decreased by 0.49 ± 0.035, with FODZYME® decreasing this change by 34.7% (−0.49 to -0.32, P=0.002) **[Figure 2B]**.

**Figure 2:**
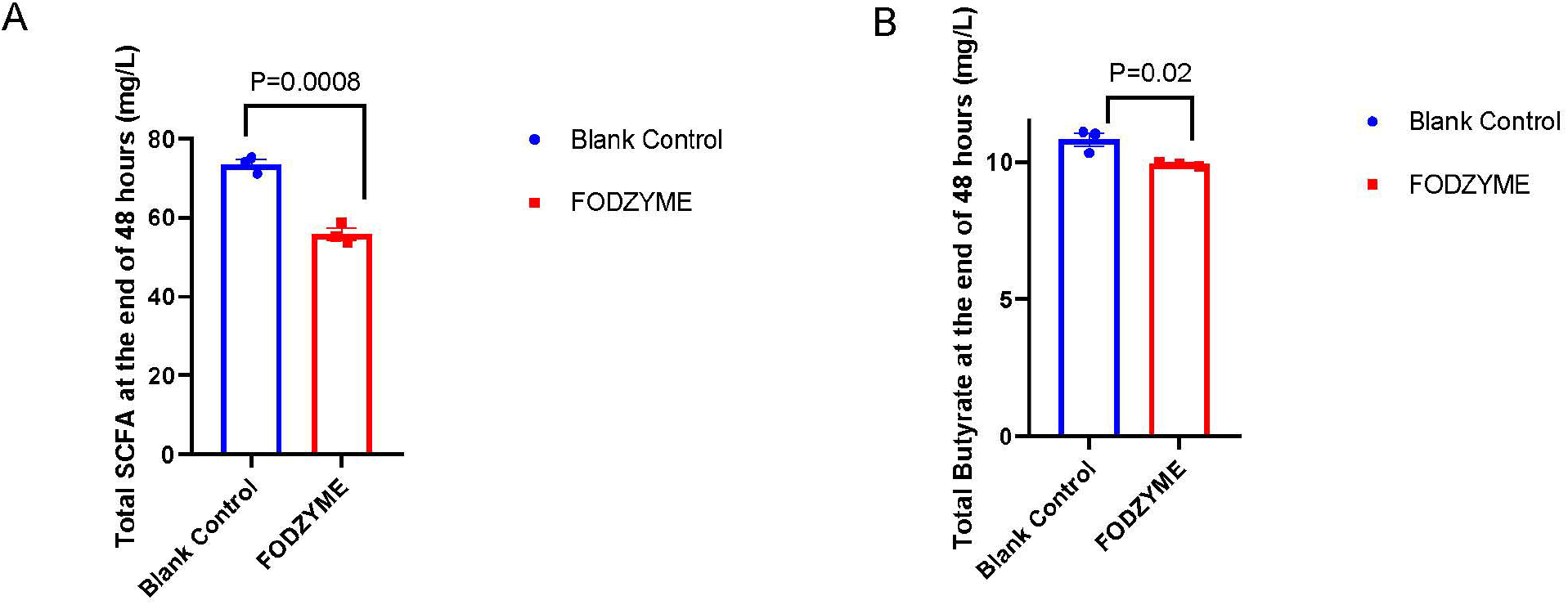
Gas Production and Change in pH in the Simulated Colon. Following 48 h of incubation in the simulated colonic environment of the SHIME®, A. gas production and B. pH change, was determined with no enzyme addition (blank control) and with FODZYME® addition. All experiments were performed in triplicate to account for variation. Error bars represent standard error.

### Short-Chain Fatty Acid (SCFA) production is preserved, with the least impact on beneficial SCFAs such as Butyrate

As one of the major concerns with a low-FODMAP diet is colonic deprivation of essential SCFAs, especially butyrate^16^, we assessed SCFA products of fermentation, noting a mild decrease in overall SCFA and butyrate production consistent with the decreased presentation of FODMAPs to the colonic environment. The observed level of total SCFA and butyrate production following FODZYME® supplementation appears to be reassuring for overall colonic health, as colonocytes depend on butyrate for up to 80% of their energy supply^22^. Compared to control conditions, total SCFA production with FODZYME® exposure at the end of 48-hour colonic simulation represented a small, but statistically significant, decrease of 19% (Control (Control 73.46 ±2.1 mM vs. FODZYME® 55.85 ± 2.6 mM, P=0.0008)) **[Figure 3A]**, with an even smaller decrease in butyrate production of 8% (Control 10.81 ± 0.42 mM vs FODZYME® 9.93 ± 0.07, P=0.02) **[Figure 3B]**.

**Figure 3:**
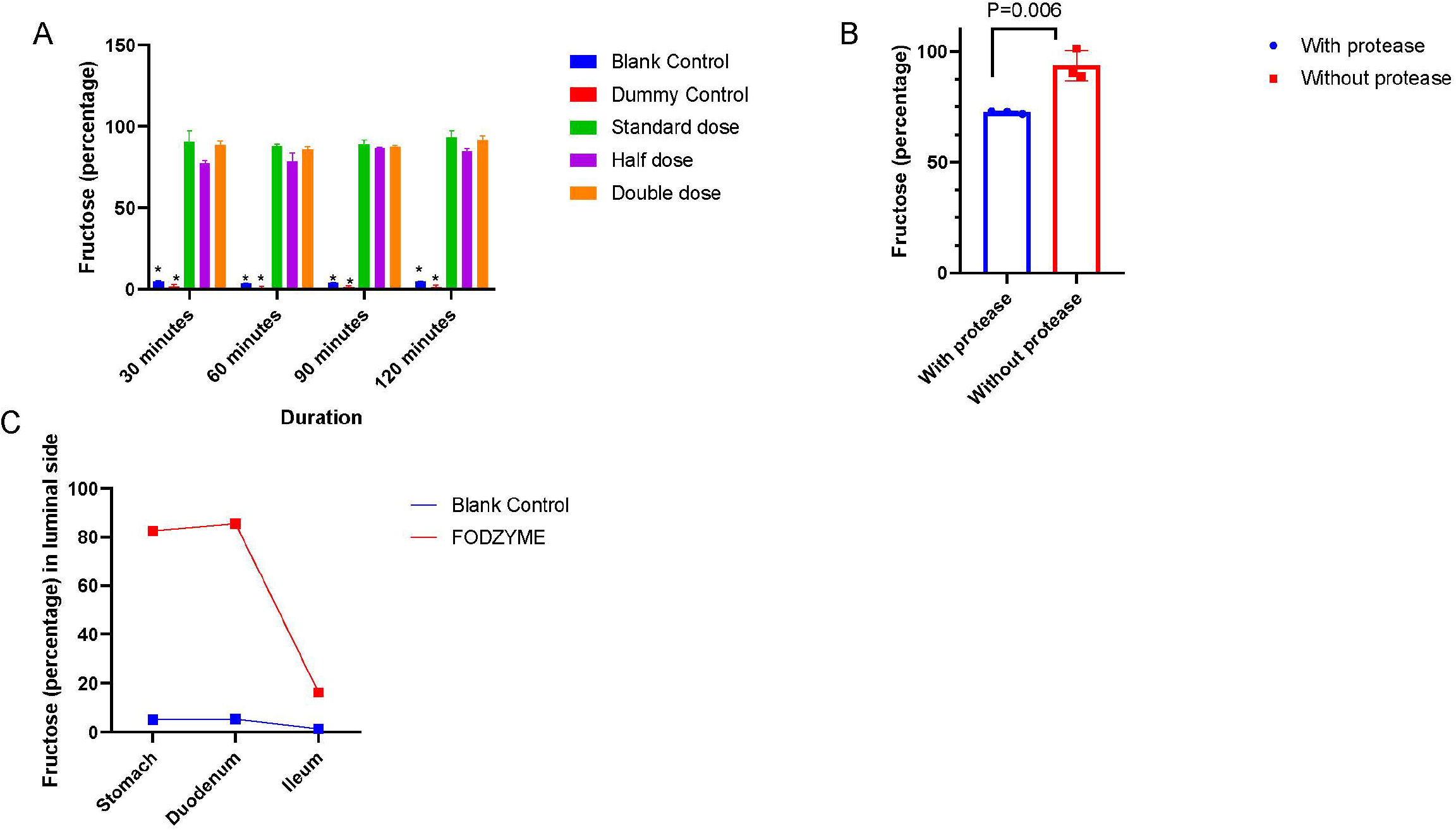
Short Chain Fatty Acid (SCFA) and Butyrate Production in the Simulated Colon. Following 48 h of incubation in the simulated colonic environment of the SHIME®, A. Total SCFA (mg/L) and B. Total butyrate (mg/L), were measured with no enzyme addition (blank control) and with FODZYME® addition. All experiments were performed in triplicate to account for variation. Error bars represent standard error.

## Discussion

FODMAPs are short-chain carbohydrates highly associated with gastrointestinal symptoms in at-risk patients such as cohorts with IBS, but also provide substrate for the production of highly important bacterial metabolites in the form of SCFAs^21^. Fructans (of which inulin represents a common source) have been increasingly identified as one of the strongest FODMAP triggers of gastrointestinal symptoms^20, 21^. Patients with IBS, a highly prevalent (11% globally^3^) condition defined by abdominal pain and alteration to intestinal function, are particularly vulnerable to symptom exacerbation from dietary FODMAPs. The varied clinical presentation of IBS often complicates management, and currently, most therapies focus on targeted symptomatic treatment^26^. Of these therapies, the low-FODMAP diet appears to have the highest efficacy, with 52-86% of patients demonstrating symptomatic improvement^4, 5, 9, 10, 17^. However, the low FODMAP diet has several limitations including being time consuming, restrictive and cumbersome. Therefore, there is a definite need for alternate approaches to mitigate FODMAP-mediated symptoms in IBS patients.

In this study, we examined the feasibility of decreasing FODMAP-induced dietary symptoms using enzymatic digestion with FODZYME®, a generally regarded as safe (GRAS) enzymatic product, primarily targeting inulin. Our study utilized SHIME®, a high-fidelity gastrointestinal simulation replicating conditions throughout the GI tract and allowing for detailed measurement of outputs such as gas production, metabolic outputs, and other data points difficult to monitor in human subject trials. Using this model, our findings broadly demonstrate plausible biological mechanisms for decreasing intestinal gas and acid production, and gastrointestinal symptoms felt to be secondary to those effects. We successfully show FODZYME® enzymatic function in gastric conditions, converting over 70% of inulin to fructose. It should be noted that patient tolerance thresholds to fructose tend to be considerably higher than tolerance thresholds to fructo-oligosaccharides (arguably an order of magnitude). Clinical responders to the low-FODMAP diet, in a blinded, randomized reintroduction trial, reported significantly worse abdominal pain in response to 0.75-1.5 g of fructan daily compared to 10.5-21 g of fructose daily^21^. The reintroduction of fructose in this study, despite being administered at 14 times the quantity of the fructan administered in its respective experimental arm, did not result in worse symptoms in a statistically significant manner compared with FODMAP elimination. For comparison, if FODZYME® were to degrade the entire fructan content of one clove of garlic (0.5 g), this would merely introduce approx. 0.5 g of extra fructose to the meal.

We demonstrate that in simulated conditions the bulk of this fructose was absorbed, and the overall decrease in FODMAP presentation to the colonic environment resulted in a substantial and significant decrease in gas and acid production. Most importantly, we were able to demonstrate substantial preservation of SCFA, most notably of butyrate, production in a simulated colonic environment populated with a healthy donor’s stool microbiota.

These findings are of particular note to patients suffering from abdominal pain and IBS, as they suggest a high likelihood of FODZYME® efficacy in replicating the impact of a low-FODMAP diet without the element of a restrictive diet, and with preservation of bacterial metabolites like butyrate which is known to be essential to the health of the colon^16^. This work demonstrates a clear mechanistic approach to mitigating the impact of high FODMAP foods in a similar fashion to lactase and sucrase replacement therapies, allowing as-needed prophylaxis of FODMAP-induced symptoms in at-risk patients. The average American diet provides 2.6 g of inulin and 2.5 g of oligofructose per individual, per day^30^. Based on our results, it is safe to speculate that even if all of this inulin or oligofructose were presented to a FODMAP-sensitive individual in a single meal, FODZYME® supplementation would be able to safely degrade the troublesome fructans, reducing the final FODMAP burden presented to the colon.

The significance of this finding is broad. Abdominal pain represents a tremendous burden to hundreds of millions of patients worldwide, and tens of millions of Americans, with tremendous associated healthcare and economic costs^2, 31^. FODZYME®, a GRAS substance of known safety and tolerability, represents a low-cost, sustainable intervention allowing for patients with these conditions to manage symptoms prophylactically. The high safety profile allows for use as sole therapy in patients with symptoms not meeting the need for daily therapeutics like tricyclic antidepressants or selective serotonin reuptake inhibitors, as well as an adjunct therapeutic for patients with higher acuity disease. This finding has tremendous implications for patients with a myriad of conditions, ranging from IBD and non-responsive celiac disease to IBS and other diseases of gut-brain interaction (functional dyspepsia, etc), and is of particular note for patients with IBS/DGBI and benign hypermobility given the high efficacy of a low-FODMAP diet in this population^32^.

Our study is limited by several factors. First, while the SHIME® system is an excellent model of the human gastrointestinal tract, efficacy in model systems does not guarantee efficacy in human subjects, and further human trial data is indicated. Our findings demonstrate the mechanistic impact on decreasing gas and acid production, but the linkage of gas and acid with symptoms in humans has not been fully elucidated. While our understanding of the pathophysiology has mechanistic support in murine models, there is only limited data in humans^33^. Finally, while multiple studies have shown the importance of SCFAs to colonic homeostasis and regulation, the exact impact of specific SCFAs and metabolomic outputs is still a topic of significant study.

In conclusion, this study demonstrates the efficacy of the FODZYME® blend in representative gastrointestinal conditions, indicating that dietary supplementation with this blend would likely decrease intestinal FODMAP burden without the concerns of a restrictive diet.

## Notes

**Financial Support:** The work herein was funded by Kiwi Biosciences.

**Disclosures:** Dr. Samant, Ms. Liu, and Mr. Hachuel are employees of Kiwi Biosciences. Dr. Wallach and Dr. Chey serve as scientific advisors for Kiwi Biosciences. Dr. Castro, Dr. Singh, Dr. Singh, Dr. Marzorati, and Ms. Duysburgh have no conflicts to report.

### Competing Interest Statement

Dr. Samant, Ms. Liu, and Mr. Hachuel are employees of Kiwi Biosciences. Dr. Wallach serves as a scientific advisor for Kiwi Biosciences. Dr. Castro, Dr. Singh, Dr. Marzorati, and Ms. Duysburgh have no conflicts to report.

### Summary of Updates

updated language describing enzyme, edits as per reviewer suggestions

